# The Polymorphism of Globin and Virus RdRP and Protein Space

**DOI:** 10.1101/2025.08.14.670433

**Authors:** Dejun Lian, Jie Lian, Qi Dong

## Abstract

The divergence of protein sequences, structures, and functions reveals Nature’s exploration of evolutionary possibilities within a given structural fold. Globins are ubiquitous across bacteria, fungi, plants, and animals; RNA-dependent RNA polymerases (RdRPs) are responsible for the replication of RNA viruses. These proteins provide excellent models for the study of protein evolution. In this study we analyzed the polymorhpisms of globins and viruses RdRPs with structure and function. We found that the protein polymorphisms are positively correlated with residue solvent accessibility (RSA). From our analysis of *protein space* of globins and virus RdRPs, we proposed that purifying selection is the predominant force of protein evolution. We also discuss the theory of last universal common ancestor (LUCA), proposed that it may be put to test.

## Introduction

The divergence of protein sequences, structures, and functions reveals Nature’s exploration of evolutionary possibilities within a given structural fold—particularly the topology of the native conformation. By analyzing what remains conserved and what changes across a protein family’s evolution, we uncover fundamental principles governing structure-function relationships. Comparative studies of highly divergent proteins sharpen our understanding of how diverse sequences can diverge and converge on similar three-dimensional architectures. Among the most illuminating examples are the globins, long regarded as a model system for probing these principles (1–3).

Globins, ubiquitous across bacteria, fungi, plants, and animals, reversibly bind oxygen via an iron-porphyrin cofactor. Beyond oxygen transport, they participate in redox catalysis and intracellular oxygen sensing. The globin family was the first in which extreme sequence divergence was observed alongside conserved tertiary structure, their structural conservation despite sequence variability highlights the evolutionary robustness of certain folds.

RNA viruses—such as hepatitis C virus (HCV), influenza, and SARS-CoV-2—exhibit rapid adaptive evolution driven by environmental pressures. Their high mutation rates (10□³–10□□ per nucleotide per replication) and short generation times facilitate extensive genetic diversification, enabling immune evasion and drug resistance. This adaptability makes RNA viruses powerful models for studying real-time molecular evolution (4). Their success hinges on genomic plasticity: unlike DNA polymerases, RNA-dependent RNA polymerases (RdRPs) lack proofreading, resulting in error-prone replication (∼10□□ errors per cycle). Despite sequence divergence, RdRPs maintain a conserved right-hand architecture (SCOP class 2.7.7.48) with catalytic aspartates coordinating metal ions essential for RNA synthesis.

Understanding sequence-function relationships is pivotal in evolutionary biology and protein engineering. *Sequence space*-the ensemble of possible protein variants——reveals how mutations influence stability and function. Maynard Smith conceptualized evolution as a traversal through this space, where functional proteins are linked via single-residue mutations (5). The “nearly neutral network” hypothesis posits evolution proceeds along paths of minimal fitness disruption, avoiding nonfunctional intermediates, maintaining functionality despite accumulating neutral mutations. Neutral drift and purifying selection shape evolutional dynamics, with most mutations being neutral or deleterious.

Advances in structural bioinformatics and deep mutational scanning now allow systematic exploration of sequence-structure-function relationships. Integrating evolutionary data with biophysical models enhances our ability to predict mutational effects, engineer proteins, and decipher evolution mechanisms. This framework elucidates how sequence space governs evolutionary trajectories, informing protein design and evolution mechanism study.

## Materials and Methods

### Sequences and Structures

2,052 sequences of globins were retrieved from PDBsum databases. Human HBA1 var are downloaded from HbVar database. A total of 12,402 sequences of HCV isolates were retrieved from the Los Alamos HCV Sequence Database, available at http://www.hcv.lanl.gov/ (6) and GenBank. A total of 107,312 sequences of IAV isolates were retrieved from Genbank, Influenza 163 Research Database (IRD) and GISAID. Approximately 30,000 sequences of SARS CoV-2 isolates which represent the entire pandemic period were retrieved from GenBank and BV-BRC databases. The Viruses RdRPs sequences were used for analysis. A listing of their accession numbers is available from the author upon request.

The structures used in this study were determined through X-ray crystallography. We used PDB code 3KMF, 3I5K, 6r65, and 7oyg structures for analysis.

### Data analysis

Multiple protein and nucleotide sequences were aligned with BioEdit and edited by hand.

In our analysis, the solvent accessible surface area (SAS) measure was used to estimate the proportion of each amino acid residue that is accessible to solvent. This was done by taking the ratio of SAS we calculated from the actual protein structure to that of the maximum exposed surface area in the fully extended conformation of the pentapeptide gly-gly-X-gly-gly, where X is the amino acid in question (10). We then normalized SAS values by the theoretical maximum SAS of each residue to obtain relevant surface accessibility (RSA), SAS are calculated using the DSSP program (http://www.cmbi.ru.nl/dssp.html).

### Statistical analysis

Data are expressed as means ± SD. Statistical analyses were performed using the Kruskal–Wallis and Mann–Whitney *U* methods. All statistical analyses were performed using SPSS version 13.0 (SPSS Inc., Chicago, IL) with additional analysis performed using Stata/MP14 (StataCorp LP). Values of *p* < 0.05 were considered significant.

### Logistic Regression, Confidence Intervals

We used the methods and model of Bustamante CD (7) in understanding how a set of predictor variables affect a dichotomous outcome variable (polymorphic or invariant), the logistic regression is an appropriate statistical model to employ.

### The correlation of residue variability with structure

The correlation was tested using Marsh L’s method (8) and Kapp OH’s method (9). We performed several tests to find structural correlates of high evolutionary rate. We used nonparametric analyses (Wilcoxon test, Spearman correlation) because neither amino acid differences nor accessibility are normally distributed. The two-sample rank-sum Wilcoxon test was used to test if groups of sites exhibited significant differences in rates of evolution. Peason’s rank correlation coefficient was calculated to test the correlation between the accessibility of residues and variability.

### The correlation of Entropy at position with structure

To measure the degree of sequence conservation, we calculated sequence entropy for each alignment position within a protein family (11–14). Information-theoretical entropy at position of protein sequences was calculated using BioEdit. Sequences were clustered before calculating to avoid the influence of redundant sequences. We used nonparametric analyses (Wilcoxon test, Spearman correlation). The two-sample rank-sum Wilcoxon test was used to test if groups of sites exhibited significant differences in protein evolution. Spearman’s rank correlation coefficient, ρ, was calculated to test the correlation between the accessibility of residues and entropy.

We used Shannon’s entropy equation which can be formulated as below:

H(l) = -Σf(b,l)log(base 2)f(b,l)

wheref(b,l)is the frequency of amino acid b(of 20) at the alignment position (19,20)

Williamson RM Groups of amino acid physicochemical properties:

20 amino acids are classified into 9 classes by their physicochemical properties as follows:(15–17)

1[V L I M], 2[F W Y], 3[ST], 4[N Q], 5[H K R], 6[D E], 7[A G], 8[P], and 9[C].

20 amino acids are further classified into 3 classes by their hydrophobicity properties as follows:(18)

Hydrophobic: F,W,Y,I,L,M,V.

Neutral: A,G,P,S,T,C.

Hydrophilic: D,E,H,K,R,N,Q.

Site entropy were calculated as follow:

H(l) = -Σf(b,l)log(base 2)f(b,l)

Where f(b,l)is the frequency of amino acid grouped using the utilization frequencies of amino acid substitution groups as estimates for their probabilities of occurrence [ the information associated with a position is expressed as H ](24)

## Results

**Figure 1.**
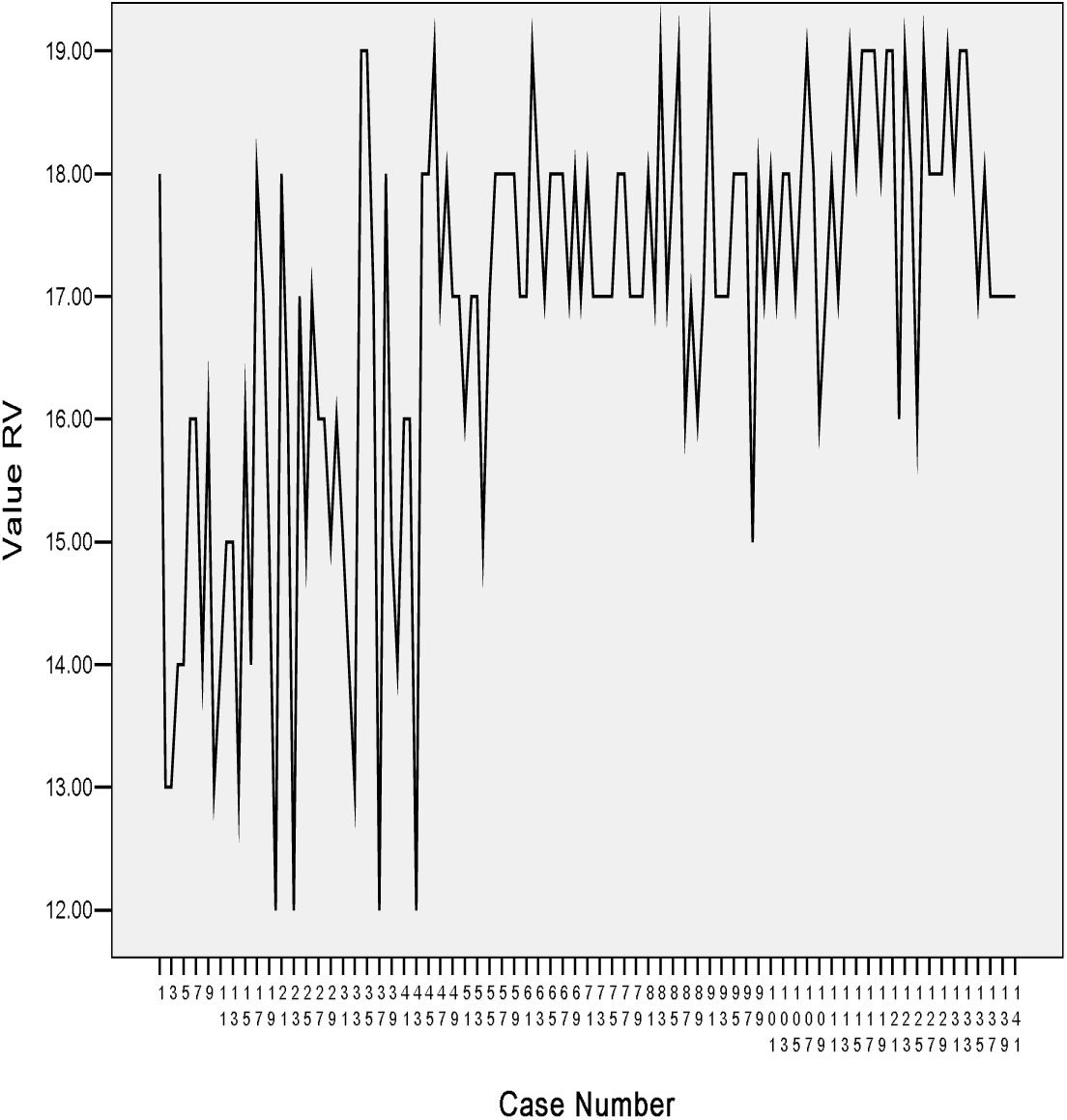
The sites polymorphism of globins.

**Figure 2.**
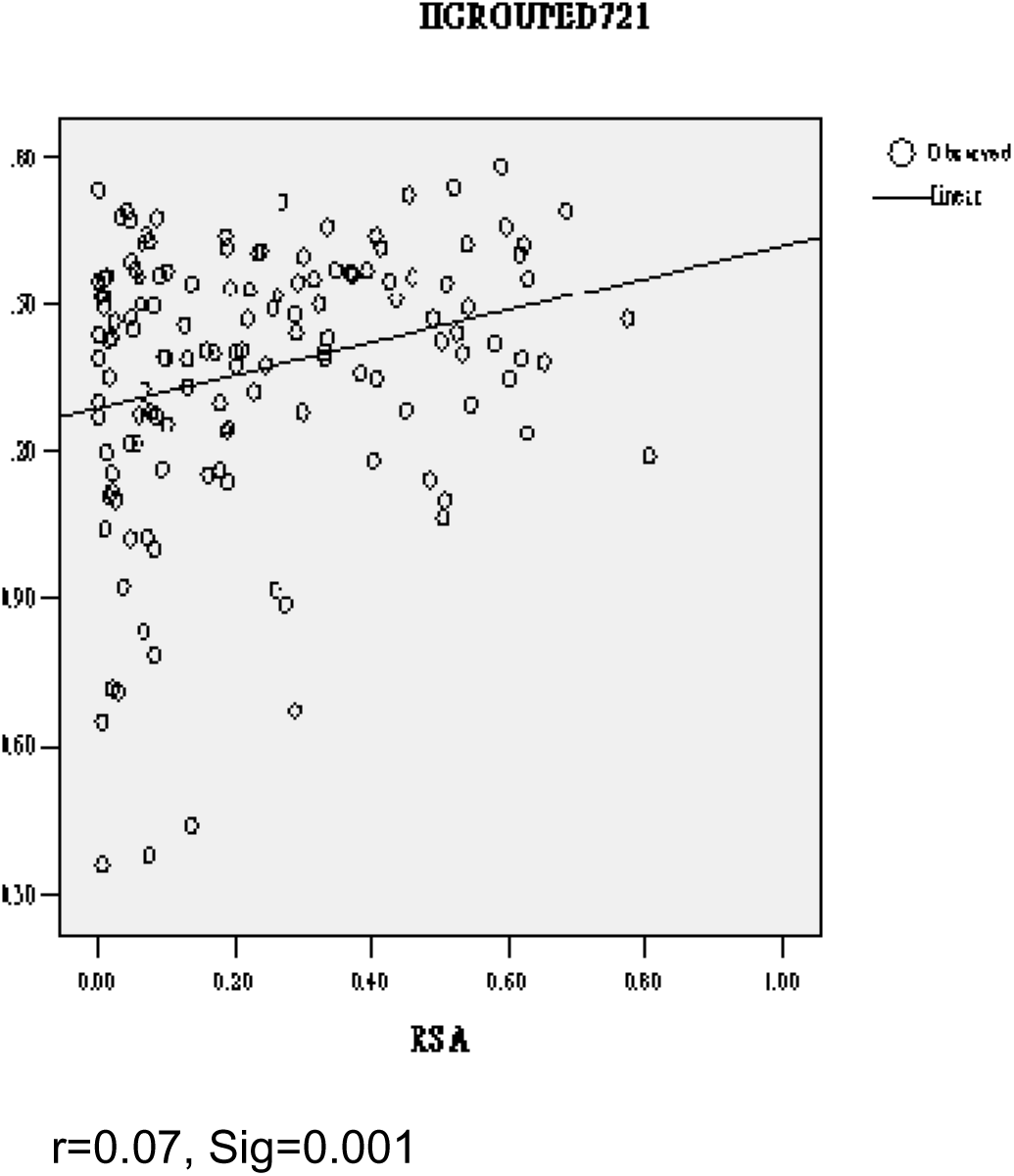
Linear correlation of residue aa grouped entropy (H) of globins with RSA

**Figure 3.**
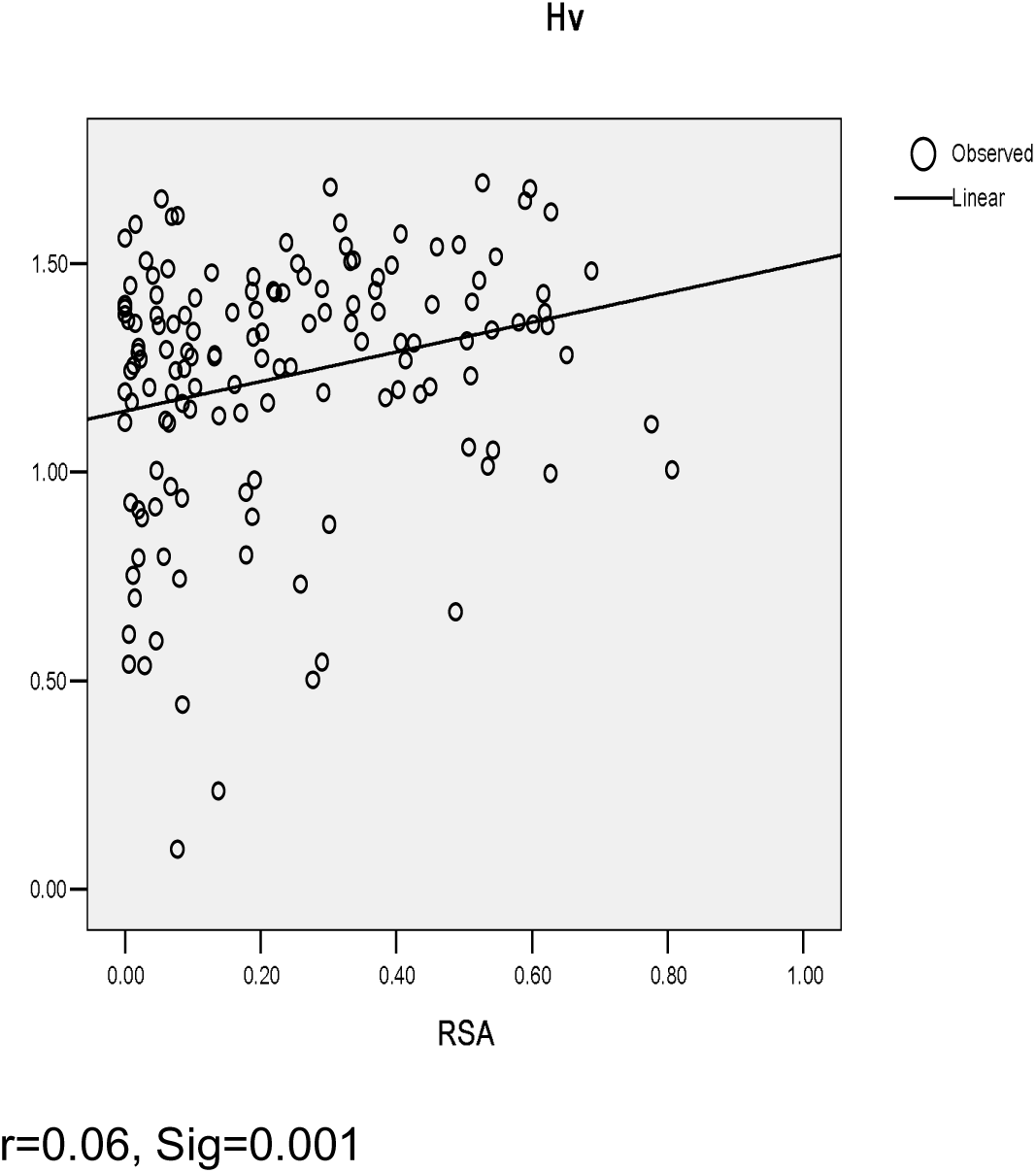
Linear correlation of residue aa entropy (H) of vertebrate globins with RSA

**Figure 4.**
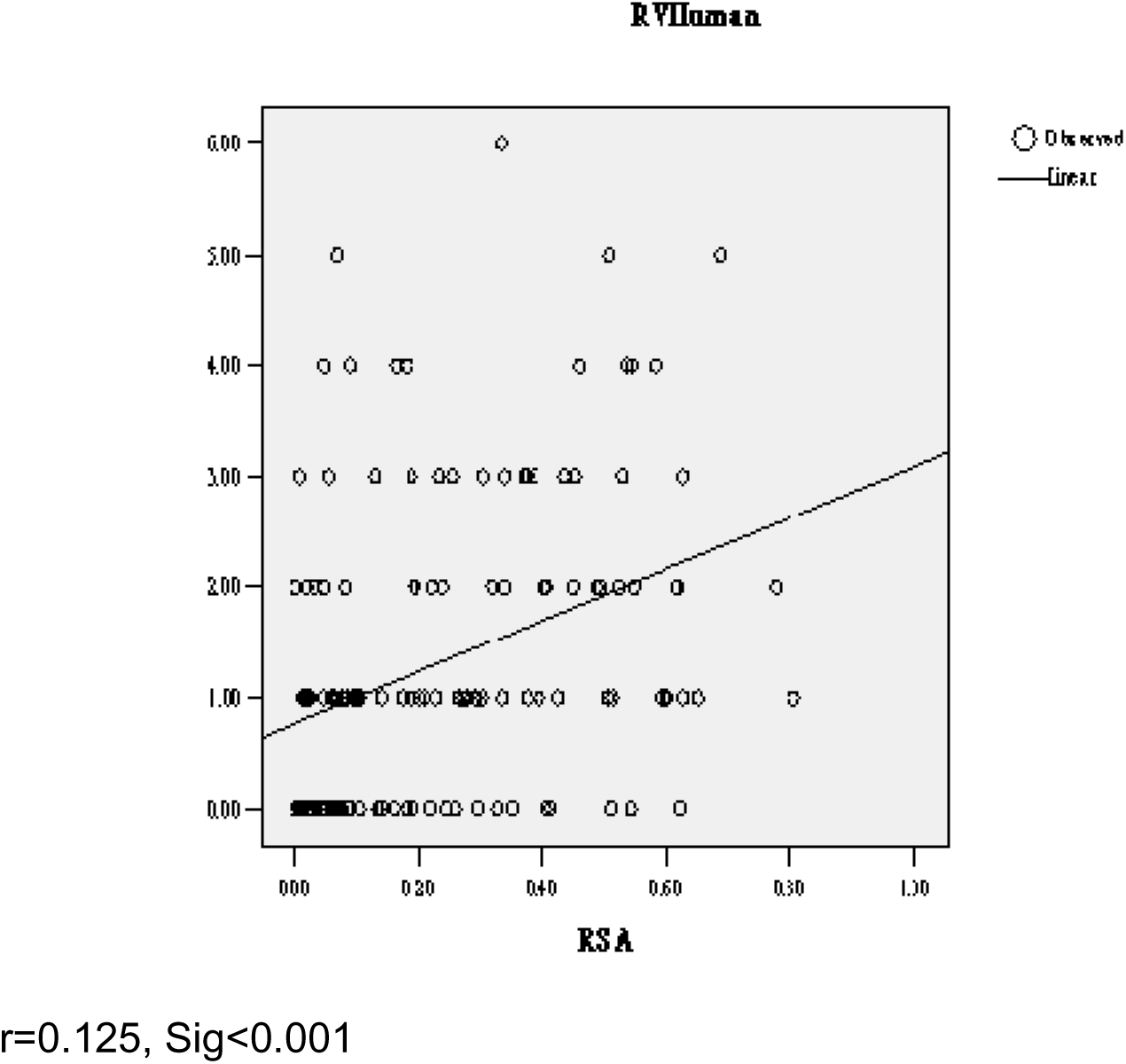
Linear correlation of RV of human hemoglobin (HBA1) with RSA

**Figure 5.**
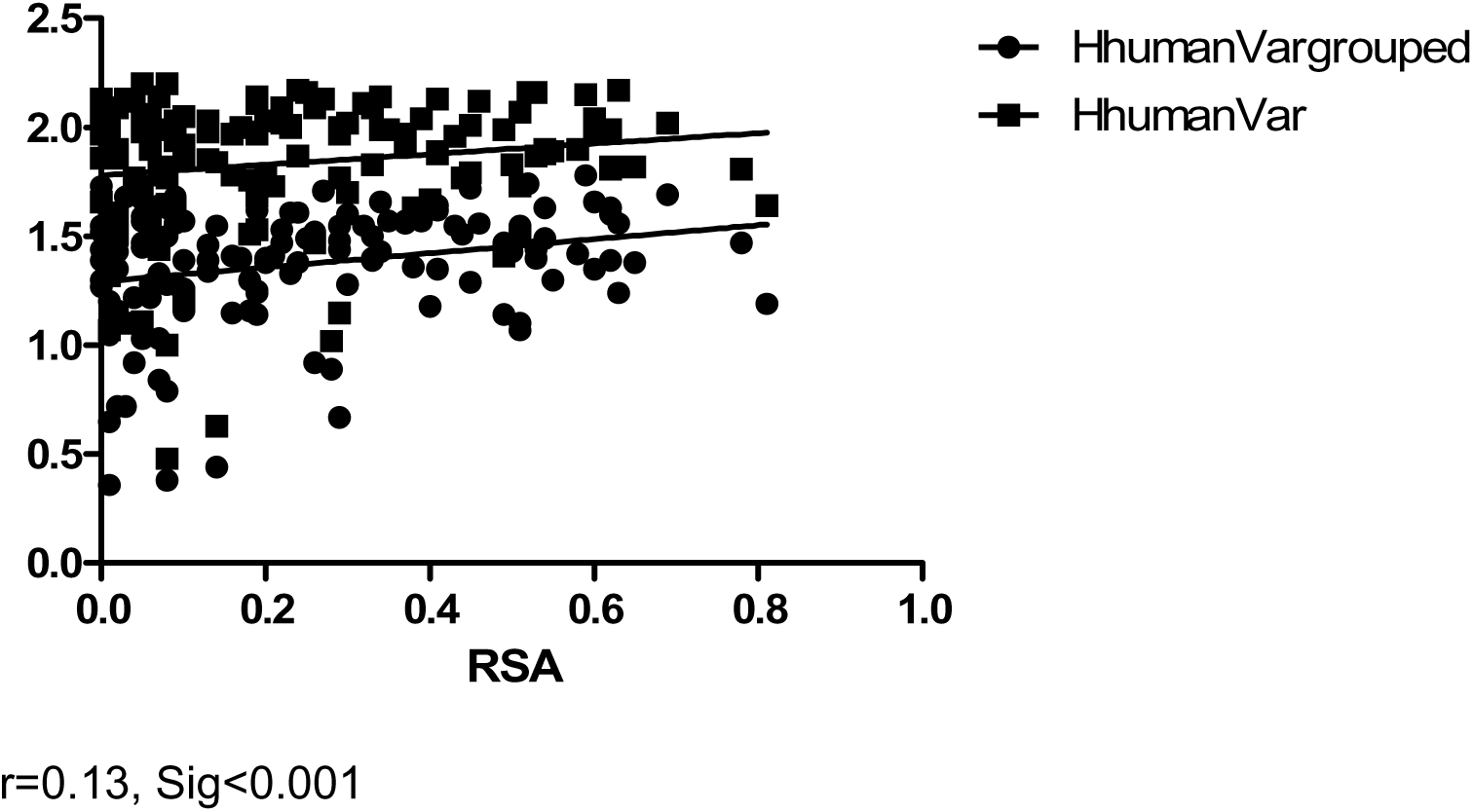
Linear correlation of Hs of human hemoglobin (HBA1) with RSA

**Figure 6.**
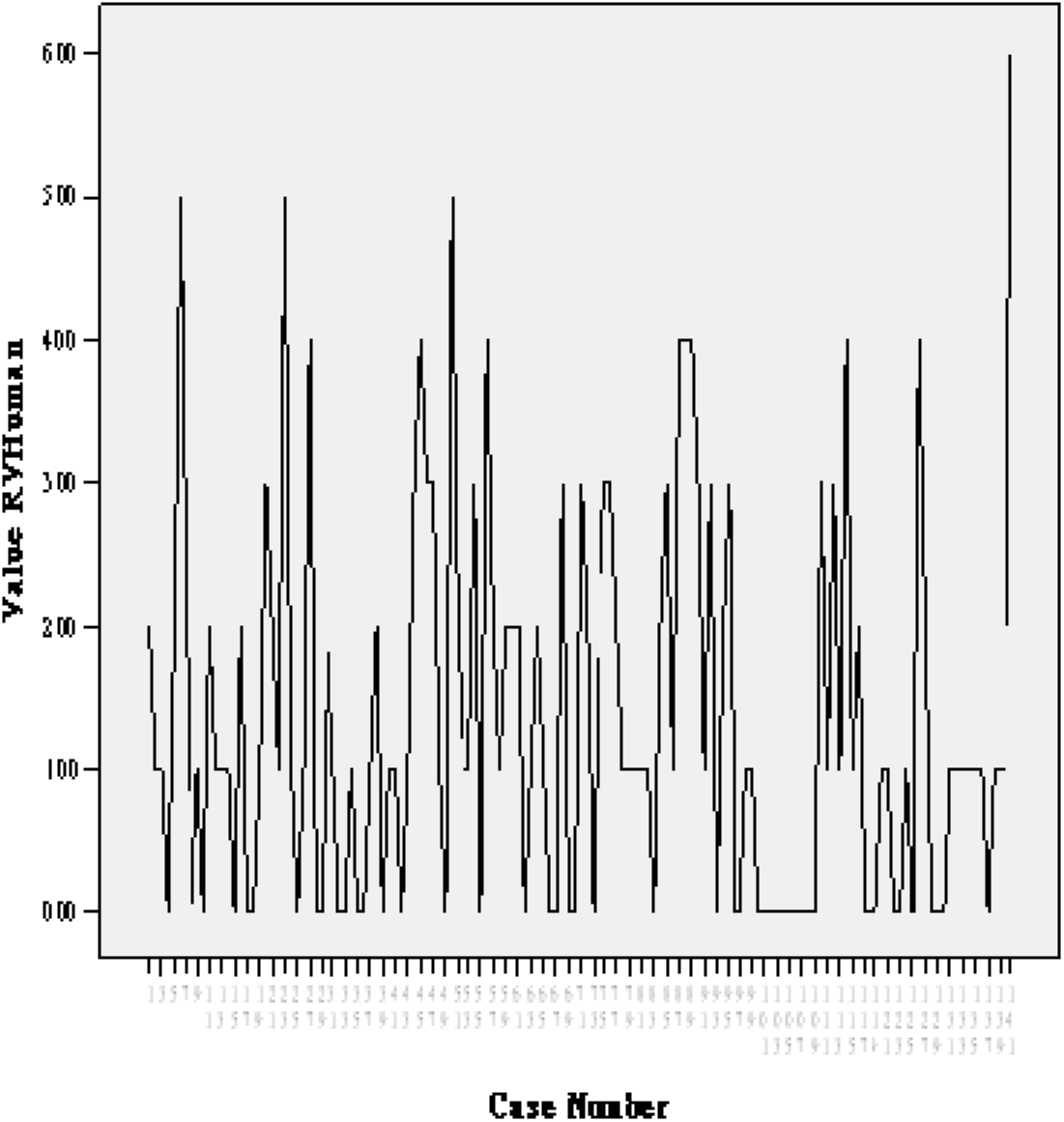
Sites polymorphism of human hemoglobin (HBA1)

**Table 1.**
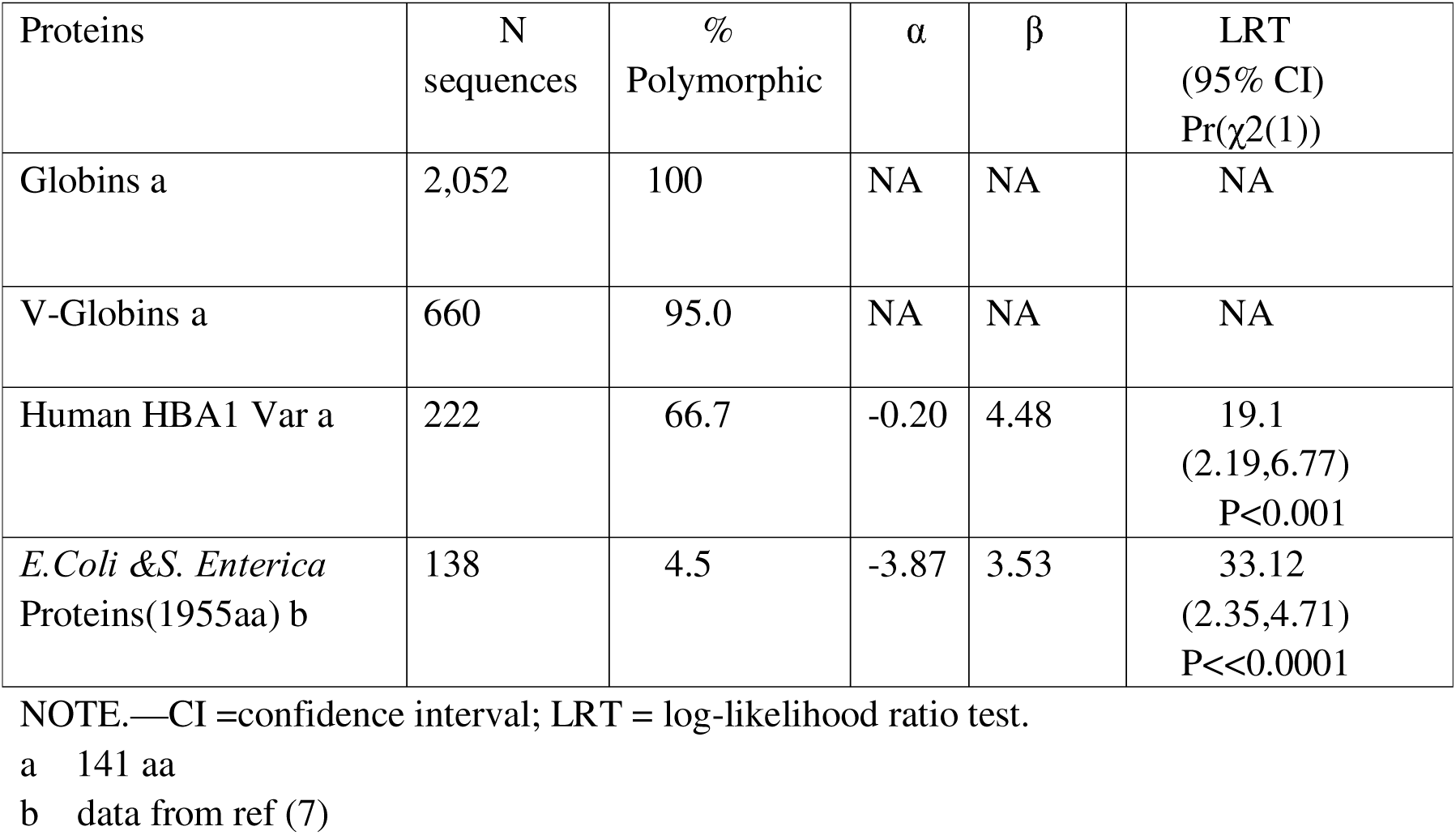
Logistic regression of globins aa polymorphism with protein structure.

**Table 2.** Logistic regression of virus RdRPs aa polymorphism with protein structure.

| Proteins | N sequences | % Polymorphic | $\alpha$ | $\beta$ | LRT (95% CI) Pr($\chi^2(1)$) |
| --- | --- | --- | --- | --- | --- |
| SARS CoV-2 RDRP | ~30,000 | 39.7 | -1.84 | 2.12 | 27.6 (1.33,2.90) P<0.001 |
| IAV PB1(757aa) | 107,312 | 95.1 | -1.17 | 2.41 | 36.71 (1.61, 3.20) P<0.001 |
| HCV NS5B (566aa) | 1,2402 | 95.2 | 0.92 | 3.52 | 32.07 (2.12, 4.92) P<0.001 |
| *E.Coli* & *S. Enterica* Proteins(1955aa) a | 138 | 4.5 | -3.87 | 3.53 | 33.12 (2.35,4.71) P<<0.0001 |

NOTE.—CI =confidence interval; LRT = log-likelihood ratio test.
a data from ref(7)

**Table 3.**
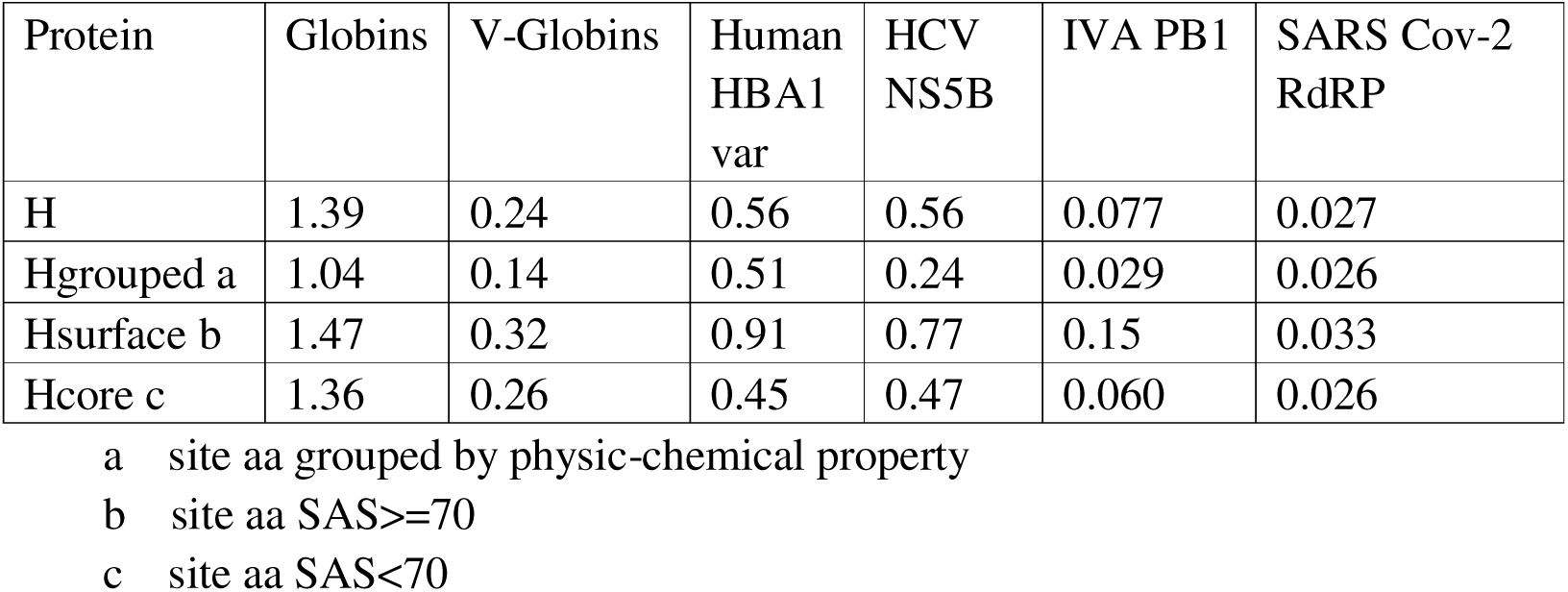
Mean entropy (H) of protein sequences.

**Table 4.** Logistic regression of human HBA1 aa grouped.

| | $\alpha$ | $\beta$ | LRT<br>(95% CI)<br>Pr($\chi^2(1)$) |
| --- | --- | --- | --- |
| Human HBA1 aa<br>grouped a | -0.83 | 2.34 | 7.83<br>(0.64,4.02)<br>P=0.006 |
| Human HBA1 RV b | -1.06 | 1.39 | 15.1<br>(0.64,2.12)<br>P<0.001 |
| Human HBA1 H c | -1.11 | 0.63 | 20.9<br>(0.33,0.94)<br>P<0.001 |

- a logistic regression of sites aa grouped by hydrophobicity with RSA b logistic regression of aa grouped by hydrophobicity with site RV c logistic regression of aa grouped by hydrophobicity with site entropy

**Table 5.** Correlation of aa polymorphism of globins with RSA.

| Globins | RV<br>(globins) | RV<br>(V-globins) | RV<br>(HBA1) | H | H(V-glo<br>bins) | H(HBA1) | Hgrouped | Hgrouped<br>(V-globin<br>s) | Hgrouped<br>(HBA1) |
| --- | --- | --- | --- | --- | --- | --- | --- | --- | --- |
| Pearson<br>corelation | 0.00 | 0.00 | 0.353 | 0.232 | 0.240 | 0.357 | 0.270 | 0.275 | 0.360 |

**Figure 7.**
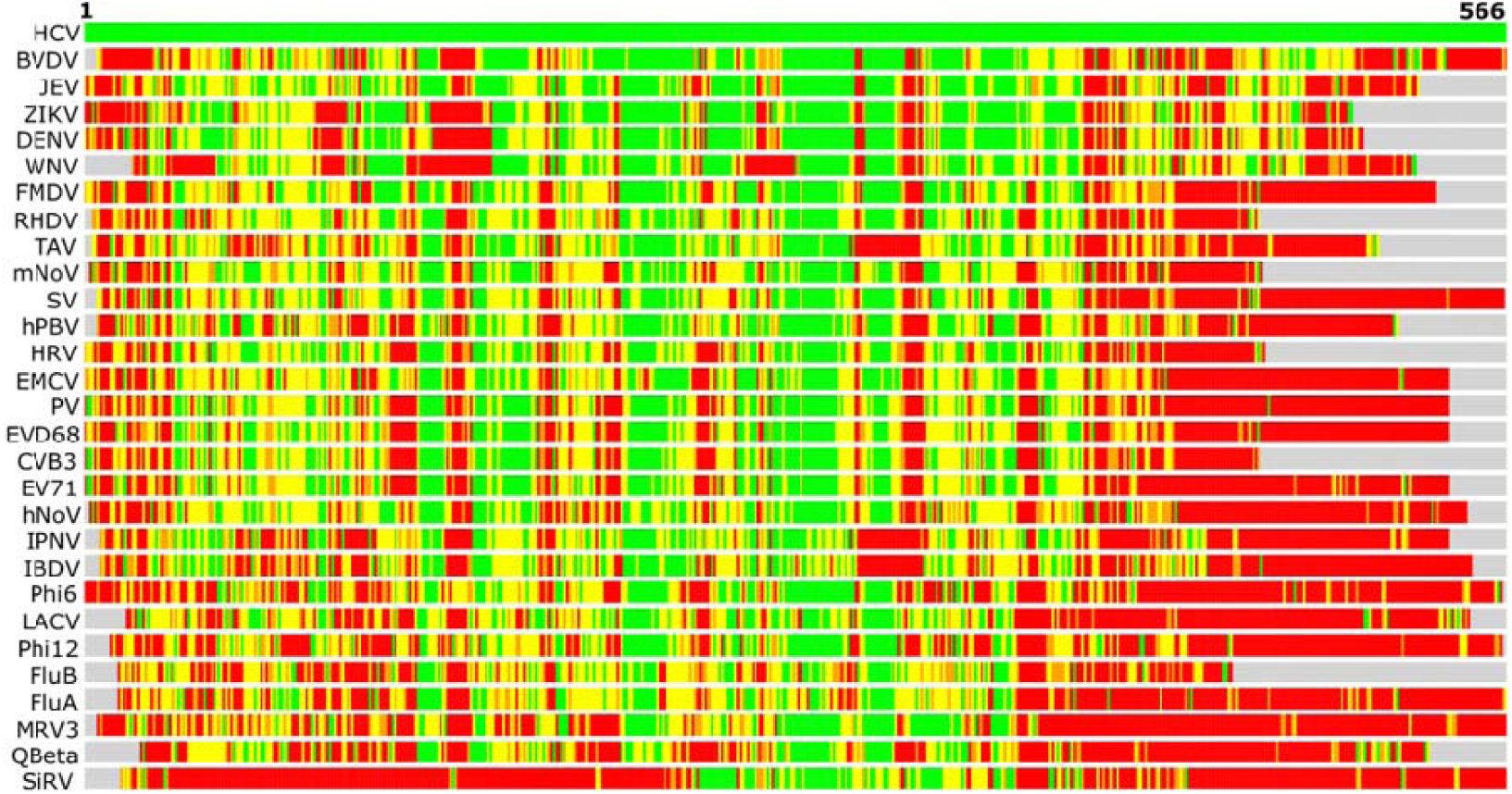
Structure-based alignment and phylogeny of representative RdRPs STRALCP server run with default parameters : (A) Structural alignment of the RdRPs of BVDV (PDB ID: 2cjq), West nile virus (WNV) (PDB ID: 2hcn), DENV (PDB ID: 4hhj), Zika virus (ZIKV) (PDB ID: 5wz3) [78], JEV (PDB ID: 4k6m), Thosea asigna virus (TAV) (PDB ID: 4xhi), Coxsackievirus B3 (CVB3) (PDB ID: 4zpc) [70], Human rhinovirus 16 (HRV) (PDB ID: 1xr7), PV (PDB ID: 1ra6) [23], Foot-and-mouth disease virus (FMDV) (PDB ID: 1u09), Encephalomyocarditis virus 1 (EMCV) (PDB ID: 4nyz), Enterovirus A71 (EV71) (PDB ID: 5f8n), Enterovirus D68 (EVD68) (PDB ID: 5xe0), Murine Norovirus (mNoV) (PDB ID: 3uqs), hNoV (PDB ID: 4nrt), Sapporo virus (SV) (PDB ID: 2uut), Rabbit hemorrhagic disease virus (RHDV) (PDB ID: 1khw), FluA (PDB ID: 5m3h), FluB (PDB ID: 4wrt) [50], LACV (PDB ID: 5amq), Qβ (PDB ID: 3mmp) [28], φ6 (PDB ID: 1hhs), Pseudomonas phage φ12 (φ12) (PDB ID: 4gzk), MRV3 (PDB ID: 1n35), SiRV (PDB ID: 2r7r), IBDV (PDB ID: 2pus), Infectious pancreatic necrosis virus (IPNV) (PDB ID: 2yi9), and Human picobirnavirus (hPBV) (PDB ID: 5i61) with HCV RdRp (PDB ID: 1nb4). The colored bars show Cα–Cα distances at each position from the amino-terminal (left) to the carboxy-terminal end (right) between HCV (top bar) and other structures. The colors indicate distances between aligned residues ranging from green (below 2 Å), yellow (below 4 Å), orange (below 6 Å), to red (above 6 Å). Figure adopted from ref(19).

The results of correlation of polymorphisms with structure and function of RdRPs of HCV, IVA, and SARS CoV-2 can be found in ref(20–22).

## Discussion

Protein evolution proceeds slowly because most amino acid substitutions impair protein structure, expression, stability, interactions, or function. These selective constraints are exceptionally stringent, as demonstrated by the structural conservation of proteins across distantly related species despite billions years of divergence and convergence evolution. Although insertions, deletions, and substitutions can drastically alter sequences during evolution, core three-dimensional structures and functions remain conserved within protein families.

Natural protein sequences emerge from a delicate equilibrium of functional, stability, and kinetic constraints. While most random mutations do not improve stability or function, neutral (or near-neutral) variants may persist under natural selection. Thus, the sequence space compatible with a specific protein fold is vast—though negligible compared to the theoretical maximum (20^N^ for an N-residue protein) (5). Estimating the size of *protein space*——a longstanding challenge in evolutionary biology——reveals its paradoxical nature: through enormous (∼10^130^ for a 100-residue protein), the fraction encoding functional folds is far small (as in the cases of globins and RNA viruses RdRPs families).

Protein evolution reflects nature’s exploration of structural and functional possibilities within a given fold. By comparing conserved and divergent features across protein families, we uncover fundamental principles governing structure-function relationships. Globins, for instance, have long served as a model system for such investigations due to their striking structural conservation despite extreme sequence divergence—a phenomenon termed “structural inertia”, now recognizes as a general principle in molecular evolution.

Globins, found in bacteria, fungi, plants, and animals, exhibit striking structural conservation despite extensive sequence divergence. Our analysis of 2,052 globins confirms this trend: low sequence homology coexists with preserved architecture. The globin fold’s resilience to sequence diversification suggests minimal structural variation (Figure 7) among the >261,309 sequenced globins. Early crystallographic studies of sperm whale myoglobin and horse hemoglobin first revealed this structural conservation amid sequence divergence (23). Subsequent work on distantly related globins demonstrated that tertiary structure persists even when primary sequences share negligible similarity. Comparative analyses (1–3) further elucidated the mechanistic basis of this inertia, a phenomenon now recognized in other protein families.

RNA-dependent RNA polymerases (RdRPs) are multidomain (α/β) enzymes that catalyze template-directed RNA synthesis using divalent metal ions [4]. The average length of the core RdRPs domain is less than 500 amino acids and is folded into three subdomains, viz., thumb, palm, and fingers resembling a right-handed cup [4]. The active sites of RdRPs from different RNA viruses are conserved and show resemblances to those of other enzymes such as reverse transcriptases and DNA polymerases, reflecting conserved nucleotidyl transfer mechanisms; but the sequences of these proteins show great diversity within and between species(24).

The protein space of globins is very vast, as results of our analysis shown, all of the 141 sites of the sequences of hemoglobin-like fold are polymorphic (Figure 1), and for all the sites the residue variability (RV) are larger or equal to 12, even the human hemoglobin α subunit (HBA1), a within species protein subunit, the maximum RV is 6. These all mean that a protein family with the same fold can adopts a large quantity of sequences of protein space.

Although the sequence space of protein families is very vast (25,26), it is not unrestrained. Mutation studies have showed that for most of sites of a protein, nearly all of the 20 amino acids can be replaced without significant influence of the function and structure of the protein (27–32). While in biological systems, this is not the case, there is very strong constraints of residue variability at nearly all sites, which shows that many of the replacement may be deleterious or slightly deleterious and will not be allowed. Mutations occur at an approximately constant rate. They can be fixed stochastically (by random drift), eliminated by negative or “purifying” selection, or they can be positively selected. Following their appearance, most mutations are purged while some are fixed not only by selection, but also by chance (33). For globin sequences analyzed, of only ∼12% of globin sites exhibit maximum variability (RV=19) while ∼30% show limited diversity (RV<=17). And for within species proteins, such as human hemoglobin α subunit (HBA1), 37 sites (36.7%) are monomorphic; and for HCV, a virus with a high mutation rate and huge population size, its NS5N protein, of the 591amino acids of this protein, 28 (4.7%) sites are monomorphic, its mean RV is 4.4. These mean that for a protein family, most of the sites are under physicochemical and physiological constraint and its protein space is limited.

The high mutation rate, huge population size, and vast sequence data of viruses provide excellent models for studying real-time molecular evolution. The results of logistic regression and liner regression analyses show that the amino acid polymorphisms of RdRPs of HCV, IVA and SARS CoV-2 are all positively correlated with RSA, as is shown that their residue variability (RV) and amino acid (aa) entropy are positively correlate with RSA (20–22), which means that the outer side of the molecular is less conservative compare with the core. The parameters of logistic regression of SARS CoV-2 RdRP are nearly the same with IVA PB1, and different with HCV and proteins of *E. coli* and *S. enteric*, which may be attributed to the fact that the evolution times of both SARS CoV-2 and IVA are short, compare with HCV and *E. coli* and *S. enteric*. The parameter β of SARS CoV-2 RdRP and IVA PB1 are smaller compare with HCV and *E. coli* and *S. enteric*, reflecting that the inner-out protein structure restriction is relaxed, so there are more polymorphism of aa of these proteins at both the surface and core. Unexpectedly, for SARS CoV-2 RdRP, the aa entropy and grouped by aa physicochemical property entropy are nearly same, which is different compare with the results of HCV and IVA PB1 (20,21). This finding can be attributed to that the evolution of SARS CoV-2 is within a short evolution period (only less than six years), and within short evolution time, the accumulated nucleic acid (na) mutants are all single mutant within coding triplet; and for single mutants of triplets, the accumulation of coding aa are random, depleted with physicochemical property restriction for aa which is observed in previous studies of protein evolution and virus evolution. These findings also mean that these may be common phenomena of the mechanism of protein evolution and virus evolution.

The Wilcoxon rank-sum test revealed significant aa entropy differences between globins and viruses RdRPs proteins surface with core regions. This divergence indicates that reduced purifying selection at solvent-accessible surfaces allows residues in high-accessibility regions to evolve at higher rate comparable to those in constrained low-accessibility zones.

In this study we used sequence entropy (H) as a measure of evolutionary conservation in a protein family. Sequence entropy served as our metric for evolutionary conservation. While valuable, entropy calculations from multiple sequence alignments may introduce bias (11). Residues were classified as either invariant (Type I) or physicochemically conserved substitutions (Type II). Following Shen & Vihinen (13) and Mirny et al. (58,59), we employed grouped amino acid entropy to analyze Type II conservation. Notably, HCV NS5B, IVA PB1, and SARS CoV-2 RdRP Shannon entropies strongly correlated with relative solvent accessibility (RSA) (20–22), confirming lower conservation at molecular surfaces. Further more, we found that their grouped Shannon entropies reduced dramatically compare with sequence entropies(Figure 5, ref 20-22), which shows conservative within aa physicochemical properties and purifying selection at aa physicochemical property level.

Studies on protein families have consistently demonstrated that amino acid substitutions occur more frequently between physicochemically similar residues ( amino acids within the same group of physicochemical property) than between dissimilar ones. We found this kind of purifying selection for globins and virus RdRPs proteins polymorphism act at conservative amino acid level –constraints on sequence diversity; while aa are grouped by their hydrophobicity, both hydrophobic and hydrophilic residues can tolerate neutral aa. This phenomenon reflects the action of purifying selection at aa physicochemical property level, a fundamental evolutionary mechanism that constrains sequence diversity by preferentially eliminating mutations that disrupt protein structure or function. At the molecular level, this selective pressure operates most strongly at conserved amino acid positions, where even minor alterations may impair protein stability, folding, or interactions. For example, substitutions that introduce drastic changes in side-chain volume, charge, or hydrophobicity are more likely to be deleterious and thus removed from the population. Conversely, mutations preserving key physicochemical properties are more likely to be tolerated, leading to the observed bias toward conservative replacements. This “structural and functional constraint” explains why certain protein domains—particularly those involved in catalytic activity, ligand binding; or structural integrity, structural core, and even surface—exhibit extreme sequence conservation across divergent species. So the 20 amino acids of protein space can be classed into 9 groups or 2 groups (95,96)

While tertiary structure often tolerates sequence variation, viruses RdRPs exhibit exceptional within-species conservation compared to cross-species families (e.g., globins). Only 28/591 residues of HCV BS5B were invariant, including functionally critical motifs like the GDD triad – though even these may evolve under prolonged selection (35). Core regions showed mean variability of 3.8 (vs.17 in globins), with surface residues displaying bimodal constraint patterns: some were hypervariable while others remained monomorphic, consistent with mutagenesis studies.

For decades, rates of protein evolution have been interpreted in terms of the vague concept of “functional importance”. Slowly evolving proteins or sites within proteins were assumed to be more functionally important and thus subject to stronger selection pressure. More recently, biophysical models of protein evolution, which combine evolutionary theory with protein biophysics, have completely revolutionized our view of the forces that shape sequence divergence. Slowly evolving proteins have been found to evolve slowly because of selection against toxic misfolding and misinteractions, linking their rate of evolution primarily to their abundance. Similarly, most slowly evolving sites in proteins are not directly involved in function, but mutating them has large impacts on protein structure and stability. Traditional “functional importance” models have been supplanted by biophysical frameworks where “Biophysical Constraints Shape Evolution” 1) Slow evolution reflects selection against misfolding/aggregation. 2) Most conserved sites maintain structural integrity rather than direct function. For HCV NS5B, IVA PB1, and SARS CoV-2 RdRP, surface conservation likely reflects protein-protein interaction constraints with host factors and replication complex components. It can be concluded that protein evolution reflects competing pressures: 1) Structural constraints (particularly RSA) govern baseline variability. 2) Functional interactions impose additional surface conservation. 3) Emerging viruses exhibit transient relaxation of constraints. These principles unify observations across protein families and evolutionary timescales.

The studies on the evolution of protein families with their tertiary structures show that because many differing amino acid sequences are able to attain the same three-dimensional arrangement of the polypeptide main chain, one may conclude that there may be little restraint on amino acid sequences over long periods of time. For HCV NS5B, IVA PB1, and SARS CoV-2 RdRP, as within species protein families with such high RNA mutate rate, the situation is different. Despite their great diversity, the amino acid sequences of these proteins are very conservative compare with those between species protein families such as globins and rhodopsins. DuBose RF et al’s work on the polymorphisms of Bacterial Alkaline Phosphatase of *Enteric Bacteria* showed that most of replacement was conservative and the DNA sequence of silent sites were saturated (70). Sawyer RT and Hartl DL’s work on Drosophila using a Poisson random field model showed a very low value of the replacement mutation rate, relative to the silent mutation rate, These may reflect mutations occurring in only a small number of codons at which a favorable amino acid change is possible, with all other changes being strongly detrimental (71). So that there is a strong conservatism of amino acids of proteins within species.

For Globins, all the 141 sites of HBA1 fold are polymeric, even the active sites Phe 43 and His87 are polymorphric. For HCV NS5B, of the 591amino acids of this protein, only 28 aa are conservative, other 563 aa are polymorphic; many functional residues like triad GDD are polymorphic. IVA PB1 and SARS CoV-2 RdRP, have the same situation, their active site XDD are polymorphic. As suggested by Wellner A’s result, the laboratory enabled active site K219S exchange of PGK, which suggested that given enough time and variability in selection levels, even utterly conserved and functionally essential residues may change (52).

Although there is considerable variability of the inner residues (maximum residue variability, 11), the inner part of HCV NS5B is very conservative with a mean of residue variability 3.7+_0.15 (compare with globin family with 17.0+-0.16), and 15 sites are monomorphic. For IVA PB1 and SARS CoV-2 RdRP, the situation are nearly the same. These reflect strong constraint at the structure core. This structure core conservation can be attributed to that these residues are not directly involved in function but are required for maintaining three-dimensional structure, mutating them has large impacts on protein structure and stability (38)

Similarly, most slowly evolving sites in proteins may be necessary because of selection against toxic misfolding and misinteractions, linking their rate of evolution primarily to their abundance. Another reason may be the need to reduce the stability of incorrectly folded states and the forming of folding nucleus (34). Globin and Cyt C. for example, a helix has a hydrophobic patch that ought to pack against another surface, there should not be a competing hydrophobic patch on the wrong side of the helix. This would help to explain why the surface rules are not in effect at all surface sites, and are not very strict even where they are ineffective. Breaking up the continuity of an incorrect hydrophobic patch does not require a rule at every surface site; and a violation might not be too damaging if there are no other violations nearby (43–46).

Furthermore, part of those solvent-accessible positions appear to have very little restriction in substitution, while others have strict constrains and remain monomorphic. For HCV NS5B, the mean RV is 6.0, and there are 5 sites which are monomorphic. This strict conservation of surface residue may also reflect that there are functional constraints of this protein by the interaction with other proteins. Various studies have showed that protein structure and sequences might be further constrained by selection for interactions among proteins (92). For HCV NS5B, IVA PB1, and SARS CoV-2 RdRP, many studies have showed that they have interaction with host proteins. There may also be interaction of RdRPs which form RNA replication complex together with other nonstructural proteins and as yet unidentified host components. Many of these strictly conservative surface residues may involve in these interactions. However, most amino acid replacements with small effects are expected to be deleterious. It has been noted that the low frequencies of most amino acid polymorphisms in natural populations of *E. coli* and *S. enteric* imply that the mutations are slightly deleterious (5), and in the context of the stability-aggregation-degradation model, It is of interest that virtually all of these are physically located in regions of high solvent accessibility on the ‘‘outside’’ of the molecule.

At the molecular level, this selective pressure operates most strongly at conserved amino acid positions, where even minor alterations may impair protein stability, folding, or interactions. For example, substitutions that introduce drastic changes in side-chain volume, charge, or hydrophobicity are more likely to be deleterious and thus removed from the population. Conversely, mutations preserving key physico-chemical properties are more likely to be tolerated, leading to the observed bias toward conservative replacements. This “structural and functional constraint” explains why certain protein domains—particularly those involved in catalytic activity, ligand binding; or structural integrity structural core, and even surface—exhibit extreme sequence conservation across divergent species.

Purifying selection is not unique to HCV, IVA, and SARS CoV-2; it is a universal feature of molecular evolution, observed in all organisms from viruses to mammals. For example, studies on influenza hemagglutinin and HIV gp120 reveal similar patterns—despite high sequence plasticity, critical functional domains remain conserved due to structural and mechanistic constraints. Even in cellular proteins, such as histones or ribosomal subunits, extreme conservation reflects the indispensable roles these molecules play in fundamental biological processes. It can be concluded that the interplay between physicochemical constraints and purifying selection dictates the limits of protein sequence diversity. While mutation generates genetic variation, structural and functional necessity ensures that only a subset of changes are evolutionarily viable. Our results show that there may be strong purifying selection among most of the sites of HCV NS5B, IVA PB1, and SARS CoV-2 RdRP proteins (and other coded proteins). Despite the high mutation rate owning to its error-prone RNA polymerase, there is still considerable conservation of virus encoded RdRPs proteins (and its other encoded proteins, data not shown) among all genotypes found all over the world and there is strong conservative within every genotype and subtype, showing that the selection against deleterious mutations is strong. Through our analysis of protein space and evolution of globins and viruses protein, we propose that purifying selection is the predominant form of selection at the molecular level in protein evolution.

Exposure to solvent is one such structural property that has received attention as a correlation of molecular evolution, both at the whole-protein and residue levels. It was observed early in the history of structural biology (73) and has been repeatedly confirmed since, that residues buried in a protein’s core are more likely to remain conserved during evolution than their solvent-exposed counterparts.The evolutionary divergence of protein amino acid sequences is governed by purifying selection imposed by functional and biophysical constraints, which significantly suppresses the occurrence of amino acid substitutions. Under these constraints, different sites (residues) in protein sequences exhibit substantially varying evolutionary rates (the number of amino acid substitutions per unit of evolutionary time). This variation results in distinct site-specific conservation patterns in multiple sequence alignments of homologous proteins. Notably, only a small subset of sites are directly involved in protein function, and their high conservation stems from function-specific selection pressures. In contrast, mutations at the majority of sites indirectly influence fitness by affecting protein folding, stability, structure, interactions, or dynamic properties (64–66). Within various protein subsets, we identified patterns of relatively conserved residue distributions. The findings demonstrate that highly conserved residues are typically localized in regions where tertiary interactions occur, which also exhibit structural conservation. However, conservation patterns remained relatively weak across all studied cases, suggesting that protein folding information may rely more heavily on factors such as (1) secondary structure propensities and (2) hydrophobic interactions—structural determinants that do not require strict amino acid sequence arrangements. Overall, the study reveals that proteins with similar three-dimensional structures can achieve proper folding through the following mechanisms: (i) without relying on highly conserved residue clusters, and (ii) even in the absence of strongly conserved sequence profiles. In some cases, no obvious shared conservation pattern exists. Therefore, when a group of proteins exhibits a well-defined and consistent sequence conservation pattern, it is more likely to reflect a close evolutionary homology rather than mere structural constraints imposed by a particular folding topology (90).

This conservation of protein sequences may be a general phenomenon of within species protein families which reflects strict structural and functional constraints and other constraints within species. As in the study of human SNP. The mean residue variability 3.6 with a standard deviation of 3.1, which corr0.14--0.289, it was estimated that 50% of mutations are likely to have mild effects, such that they reduce fitness by between one one-thousandth and one-tenth. It was also inferred that 15% of new mutations are likely to have strongly deleterious effects (68–70). It was estimated that 19% and 20% of nonsynonymous mutations are neutral is remarkably consistent (although somewhat larger) than Eyre-Walker A et al.’s estimation of 23% (21–29%) inferred from a smaller datasets using folded frequency data and the hybrid approach. Yampolsky LY et al, expected a slightly larger proportion of mutations to be nearly neutral, they inferred that 19% of mutations are effectively neutral and that 14% of mutations are slightly deleterious, such that they segregate in the population at moderate frequencies, but never become fixed. The remainder of the mutations are strongly deleterious such that they contribute little to polymorphism or divergence (71, 72).

Clarke B(and also King and Jukes) proposed that at least 85-90% of the mutations causing a change in amino-acid are, on the average, selectively disadvantageous (75,76); Hartl DL had studied the polymorphism of amino acids based on frequencies alone using Poisson random field analysis which suggests that most polymorphic amino acids in the same proteins are slightly deleterious (5). For human SNPs, the difference in the number of rare vs. common alleles was used to estimate that 79–85% of amino acid-altering mutations are deleterious (93). It was estimated that the fraction of strongly deleterious mutations (which rarely become fixed) is > 70% in most species, only 10% or fewer of mutations seem to behave as slightly deleterious mutations (SDMs)(65). This maybe the case for RNA virus proteins. For HCV, IVA, and SARS CoV-2, with such high diversities, there may still be strong purifying selection among most of the sites of its coding proteins. It is estimated that only10^9^ out of 10^130^ sequences are accepted with a 100 aa protein (25); this does not mean that all of 10^9^ sequences are accepted sequences, most of the recombinant sequences are selection-against, only the aa mutation recombinants of protein space which form proper structure and function are selected. The analyses of Vila JA and Dryden DTF et al. had not consider the aa mutation recombinants, so they overestimated the value of accepted protein space (25,95).

From our analysis of protein space and protein evolution, we here propose that the last universal common ancestor theory of organism evolution maybe wrong. Last Universal Common Ancestor (LUCA) theory posits that all extant species on Earth evolved from the most recent common ancestor of all current life, believed to be a single-celled organism that lived over 3.5-4 billion years ago (87). Although this theory helps explain how diverse forms of life on Earth and has many proved evidence, it is not in accordance with the study of protein space and many evidence.

From our analysis of protein space of human hemoglobin and RNA viruses RdRPs, the protein space of a within species protein is quite limited owning to strong purifying selection and is quite stable, it has little possibility to evolve to a different species protein. Further more the genome of a species is very stable because of error-proof mechanism of DNA polymerases except RNA viruses; although insertion and deletion can accelerate evolution, they are uncommon. Even for RNA viruses, the protein space and genome are also quite limited because of purifying selection imposed by physicochemical and physiological constraint so they have no possibility to evolve to other species of virus from statistic point of view. Therefor the extant species must evolved from their own ancestors, at most akin species evolved from their own common ancestor; they could not evolve from a last universal common ancestor (88–90). This proposal need to be tested by further studies

